# Exposure to Airborne Bacteria Depends upon Vertical Stratification and Vegetation Complexity

**DOI:** 10.1101/2020.11.11.377630

**Authors:** Jake M. Robinson, Christian Cando-Dumancela, Rachael E. Antwis, Ross Cameron, Craig Liddicoat, Ravin Poudel, Philip Weinstein, Martin F. Breed

## Abstract

Exposure to biodiverse aerobiomes may support human health, but it is unclear which ecological factors influence exposure. Few studies have investigated near-surface green space aerobiome dynamics, and no studies have investigated aerobiome vertical stratification in different green spaces. We used columnar sampling and next generation sequencing of the bacterial 16S rRNA gene, combined with geospatial and network analyses to investigate aerobiome spatio-compositional dynamics. We show a strong effect of habitat on bacterial diversity and network complexity. We observed aerobiome vertical stratification and network complexity that was contingent on habitat type. Tree density, closer proximity, and canopy coverage associated with greater aerobiome alpha diversity. Grassland aerobiomes exhibited greater proportions of putative pathogens compared to scrub, and also stratified vertically. We provide new insights into the urban ecosystem with potential importance for public health, whereby the possibility of differential aerobiome exposures appears to depend on habitat type and height in the airspace.

## INTRODUCTION

Exposure to biodiverse environmental microbiomes – the diverse consortium microorganisms in a given environment – plays an important role in human health (***1–5***) From an early age, a complex network of environmental microorganisms supports the development and regulation of immunity (***6***). Indeed, exposure to a wide range of microorganisms is thought to strengthen our response to noxious stimuli (e.g., pathogens) and reduce the likelihood that our immune systems will be oversensitive to innocuous agents, such as dust particles, pollen, and our own cells–the latter manifesting as autoimmunity (***7–9***).

Urbanisation and loss of macro-biodiversity are linked to loss of microbial diversity, which could negatively impact the health-supporting microbial communities residing in and on human bodies – the human microbiome (***10, 11***). This loss of microbial diversity underpins the *biodiversity hypothesis*, which draws a link between concurrent global megatrends of biodiversity loss (including microorganisms) (***12**)* and rapid increases in noncommunicable diseases (NCDs) (***13***). A recent study empirically tested this hypothesis and found that exposure to plant diversity and associated microbial communities significantly correlated with reduced risk of acute lymphoblastic leukemia by promoting immune maturation (***14***).

Furthermore, biodiverse environments could supplement human microbiomes with functionally important microorganisms. Short chain fatty acids (SCFAs) are produced by certain bacteria as metabolic by-products and are known to play important roles in supporting human health. For example, the SCFA *butyrate* is linked to the inhibition of intestinal tumours (***15***) and atherosclerosis (***16***), as well as supporting bone formation (***17***) and promoting epithelial integrity (***18***). Such microorganisms may be transferred through aerobiomes. For example, in a randomised controlled mouse study, a putative soil-associated butyrate-producing bacteria was found to be supplemented in mice gut microbiota following trace-level airborne soil dust exposures and subsequently linked to reduced anxiety-like behaviour (***5***).

The aerobiome—the collection of microorganisms in a given airspace—is an important source of environmental microorganisms (***19–21***). Despite this importance, only limited studies have investigated the dynamics of near-surface aerobiomes in urban green spaces. Mhuireach et al. (2016) showed that aerobiomes in urban green and grey spaces had distinct compositions (***22***). Subsequent studies have shown vegetation type has a potential modulating effect on aerobiome diversity and composition (***23, 24***). Stewart et al. (2020) found that aerobiomes varied in composition and function between urban and suburban sites (***25***). Mhuireach et al. (2019) identified localised influences on aerobiomes, including weather and land management (***22, 26***). Our recent work has also demonstrated aerobiome vertical stratification between ground level and 2 m heights in an urban green space (***27***). Together, these studies suggest that individuals may be exposed to different aerobiomes depending on the type of habitat visited and human-scale height-based variation in environmental aerobiomes. Consequently, understanding the effects of habitat and height—and their interactions—on aerobiomes could have important implications for public health.

There is growing recognition that urban green spaces are important for human health and wellbeing through provision of psychosocial and biological benefits (***28–32***). Gaining a deeper understanding of urban green space aerobiome exposure potential could inform public health and environmental management strategies in the future. In this study, we used an innovative columnar sampling method to sample aerobiome bacterial communities in three urban green space habitat types in the Adelaide Parklands, South Australia. These habitats included amenity grasslands, woodland/scrub (dominated by native *Eucalyptus spp.* trees and shrubs; henceforth referred to as ‘scrub’), and bare ground habitat; each is a typical urban green space habitat. We conducted next generation sequencing of the bacterial 16S rRNA gene to characterise the diversity, composition and network complexity of aerobiomes. We also applied geospatial analytical methods to explore the potential influence of trees on the micro-biodiversity of aerobiomes. Our primary objectives were to: **(a)** assess aerobiome composition and micro-biodiversity differences between the three habitats; **(b)** compare aerobiome vertical stratification between the different habitats; **(c)** assess whether tree density, distance to trees, and tree canopy coverage influenced bacterial alpha diversity; and, **(d)** to assess any differences in known pathogenic bacterial taxa between habitats and sampling heights.

## RESULTS

Bacterial communities were dominated by three key phyla in all three habitats: Proteobacteria, Bacteroidetes, and Actinobacteria, however, abundance differed depending on height (**Fig. 1**) (full description of sequencing reads in Supplementary Materials, Appendix B). We now present the results in order of the objectives (a-d) set out in the Introduction.

**Fig. 1.**
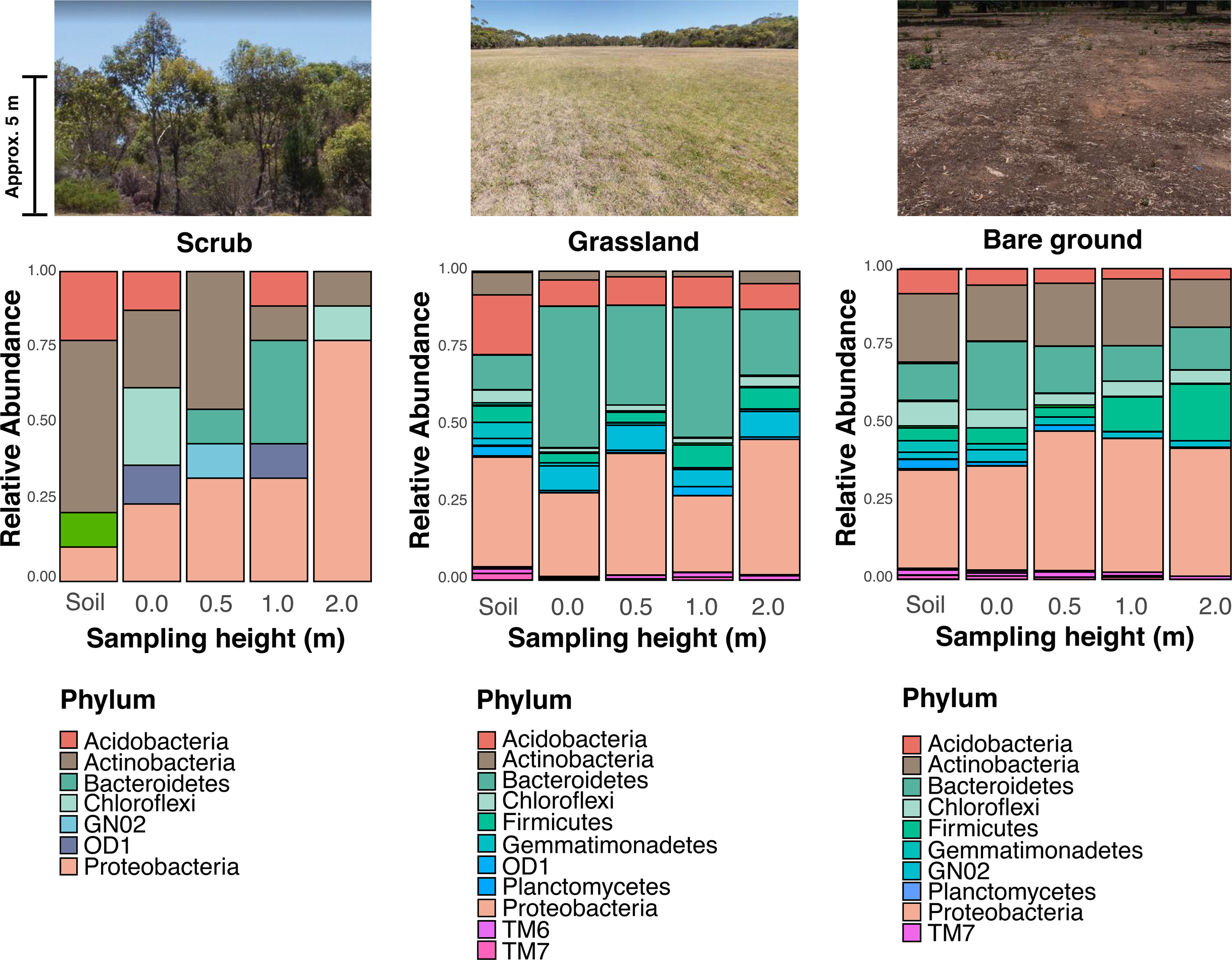
Profile of bacterial communities from each habitat at the phylum level. The coloured area of each bar represents the relative abundance of the corresponding phylum over 1%. The X-axis displays sampling heights: soil, 0.0 m, 0.5 m, 1.0 m, and 2.0 m (from left to right). The photographs above the plots show examples of each habitat used in the study (photographs by authors).

### Comparison of bacterial alpha diversity between habitats

We found that bacterial alpha diversity of the soil differed significantly between habitats (ANOVA F = 3.95, df = 1, *p* = 0.03) (**Fig. 2**). The soil microbiome from the scrub habitat was significantly more biodiverse than the grassland habitat (Tukey multiple comparison of means test; scrub x̅ = 5.78; grassland x̅ = 5.46; adjusted *p* = 0.02). We also found that bacterial alpha diversity of the air differed significantly between bare ground and scrub habitats (Chi-squared = 11.3, df = 1, *p =* <0.01), with the scrub aerobiome being more biodiverse than the bare ground. Aerobiome alpha diversity of scrub and grassland were also significantly different (Chi-squared = 24.8, df = 1, *p =* <0.01), and the scrub aerobiome was the most biodiverse. No significant difference was observed in alpha diversity between bare ground and grassland habitats (Chi-squared = 0.46, df = 1, *p =* <0.49).

**Fig. 2.**
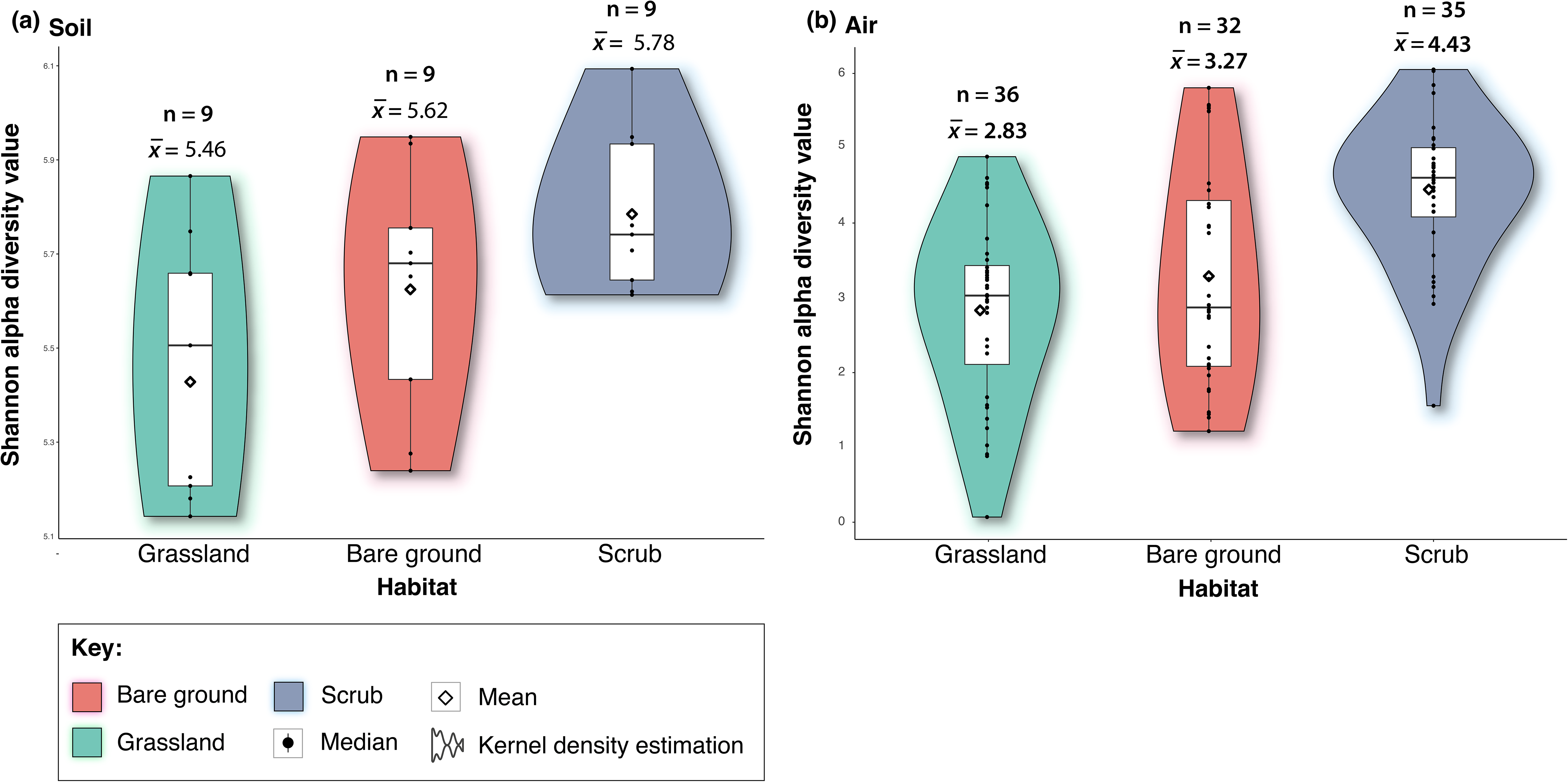
Box/violin plots of Shannon alpha diversity scores for each habitat; bare ground, grassland, and scrub. Panel (a) shows inter-habitat bacterial alpha diversity for soil samples, and panel (b) shows inter-habitat bacterial alpha diversity for aerobiome samples. Plots also display mean and median values, interquartile range and kernel density estimation (Shannon alpha diversity values for each habitat, divided into days and sites, are in Supplementary Materials, Appendix B). We also tested for mean alpha diversity differences between dates and sites, showing that sampling dates and individual sites were generally not a factor in alpha diversity variation with nearly 90% of comparisons showing non-significant results.

### Comparison of bacterial beta diversity between habitats

We observed clear differences in aerobiome compositions (beta diversity) (PERMANOVA, df = 2, F =3.7, R^2^ = 0.07, *p* = <0.01, permutations = 999) and soil samples (PERMANOVA, df = 2, F =6.8, R^2^ = 0.36, *p* = <0.01, permutations = 999 among habitats (**Fig. 3**). For air samples, all habitats displayed significantly distinct bacterial communities, where habitat type explained 7% variation in bacteria community composition. However, there was significant heterogeneity in dispersion (PERMDISP, F = 13, *p* = <0.01). For soil only, habitat type explained 36% variation in bacteria community composition, however, this increased significantly to 75% and 74% when comparing scrub to grassland and scrub to bare ground, respectively (PERMANOVA, df = 5, F =7, R^2^ = 0.75 and 0.74, *p* = <0.01). There was no significant heterogeneity in dispersion (PERMDISP, F = 2, *p* = 0.07).

**Fig. 3.**
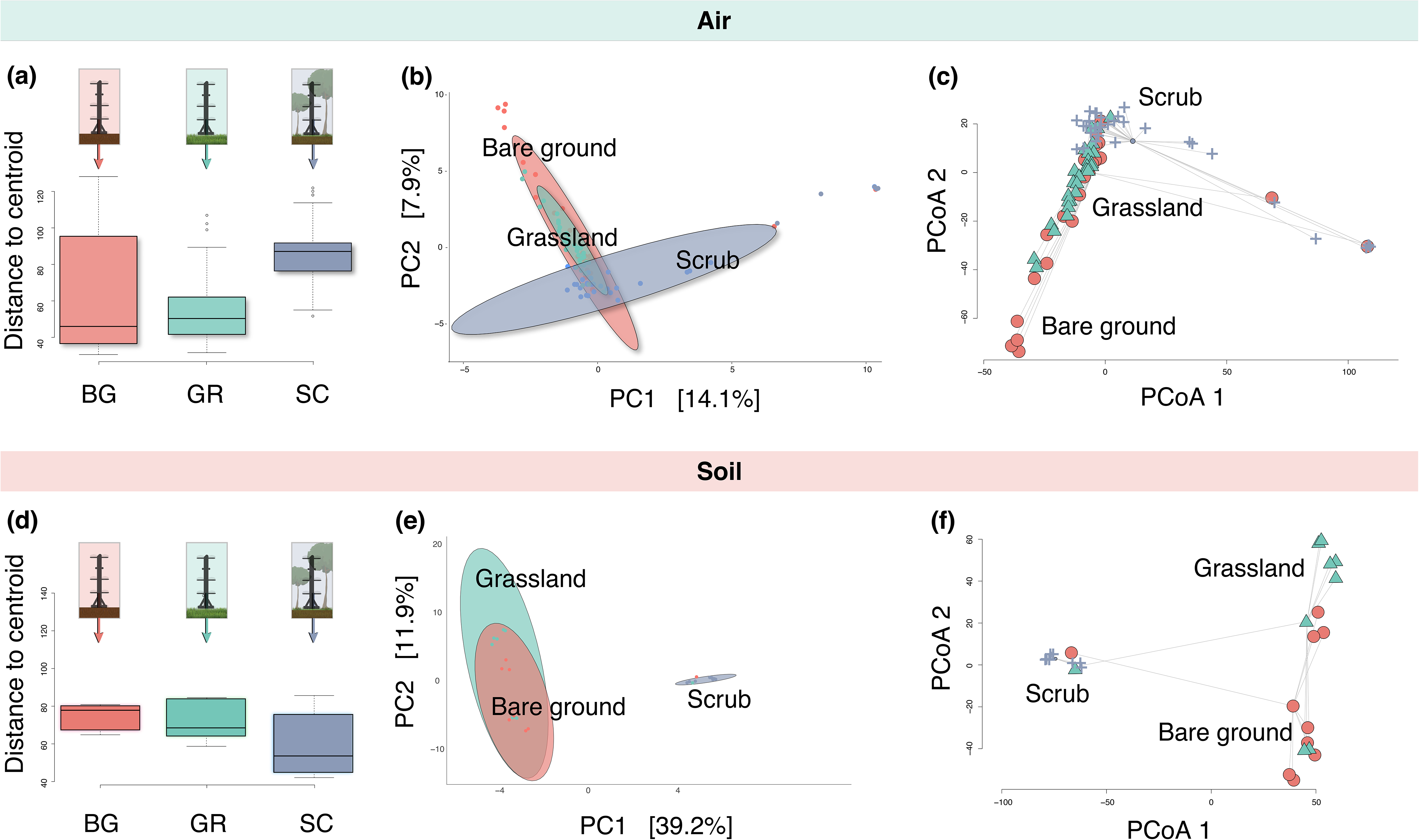
Upper panels show air samples only, whereby (a) is a boxplot of dispersion (spread); (b) ordination of bacterial communities for all habitats (BG = bare ground; GR = grassland; SC = scrub), ellipses represent Euclidian distance from the centre – with the radius equal to the confidence level (0.95); and (c) ordination of dispersion by Aitchison Distance. Lower panels show soil samples only, whereby (d) is a boxplot of dispersion; (e) ordination of bacterial communities for all habitats, ellipses represent Euclidian distance from the centre; and (f) shows an ordination of dispersion by Aitchison Distance.

### Vertical stratification: alpha diversity

#### *Bare ground v*ertical stratification: alpha diversity

For the bare ground habitat, we observed a strong negative correlation between alpha diversity (air and soil for all sites/dates) and sampling height from ground level to 2 m (Pearson’s *r* = −0.75, df = 39, *p* = <0.01) (**Fig. 4, a**). Alpha diversity (Shannon scores) ranged from 1.2 to 5.93 and was highest at soil level, followed by lower air sampling levels (0.0 m-0.5 m) and upper sampling levels (1.0 m-2.0 m), respectively. Analysis of air-only samples also showed a significant negative correlation between height and bacterial alpha diversity, demonstrating vertical stratification in this bare ground habitat (Pearson’s *r* = −0.60, df = 30 *p* = <0.01).

**Fig. 4.**
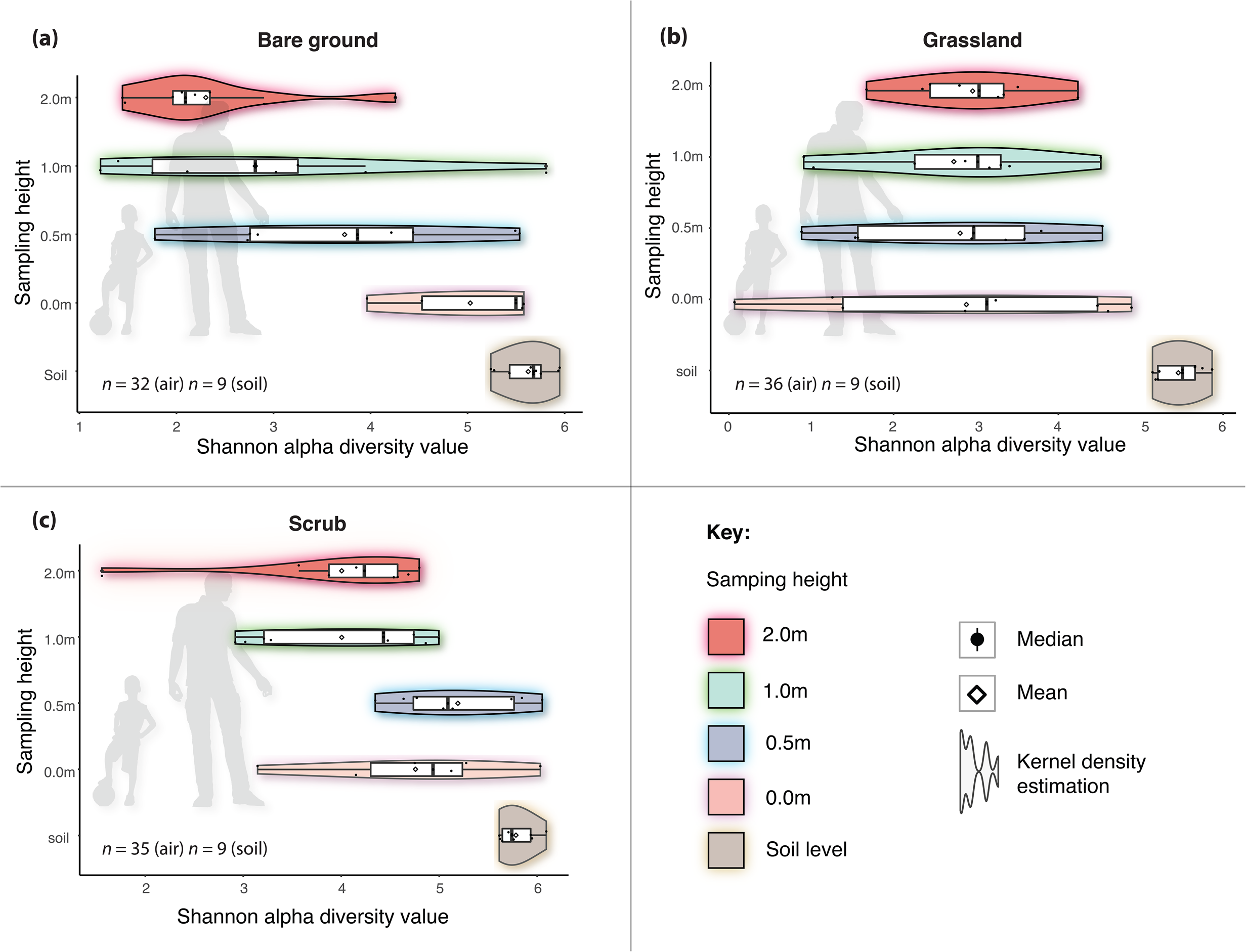
Box/violin plots of Shannon alpha diversity scores for each sampling height including soil and for each habitat: (a) bare ground; (b) grassland; and (c) scrub. Plots also display mean values, interquartile range and kernel density estimation, and silhouettes of humans for perspective.

#### *Grassland v*ertical stratification: alpha diversity

For the grassland aerobiome, we observed a significant negative correlation between alpha diversity (air and soil for all sites/dates) and sampling height from ground level to 2 m (Pearson’s *r* = −0.38, df = 43, *p* = 0.01) (**Fig. 4, b**). Alpha diversity ranged from 1.2 to 5.9 and was highest at soil level. However, once air sample data were isolated from soil sample data and analysed separately, the correlation was weak and not significant, indicating that vertical stratification was not detected in this grassland habitat (Pearson’s *r* = 0.03, df = 34, *p* = 0.86; see Supplementary Materials, Appendix B for correlations between individual dates and sites).

#### *Scrub v*ertical stratification: alpha diversity

In the scrub aerobiome, we observed a significant negative correlation between alpha diversity (air and soil for all sites/dates) and sampling height from ground level to 2 m (Pearson’s *r* = −0.59, df = 39, *p* = <0.01) (**Fig. 4, c**). Bacterial alpha diversity in the scrub habitat ranged from 1 to 6 (Shannon score) and was highest at soil level, followed by lower air sampling levels (0.0 m - 0.5 m) and upper sampling levels (1.0 m - 2.0 m), respectively. Analysis of air-only samples showed a significant negative correlation between height and bacterial alpha diversity, demonstrating vertical stratification in this scrub habitat (Pearson’s *r* = −0.38, df = 30, *p* = 0.03).

### Vertical stratification: beta diversity

#### *Bare ground v*ertical stratification: beta diversity

Sampling heights in the bare ground habitat displayed disparate bacterial compositions (**Fig. 5, a**). Sampling height explained 29% variation in bacteria community composition when all air sampling heights were included (PERMANOVA df = 4, F = 3.67, R^2^ = 0.29, *p* = <0.01, permutations = 999). Analysis of air samples for the bare ground habitat in isolation showed that sampling height still explained 25% variation in bacterial community composition (**Fig. 5, d**) (df = 3, F = 3.06, R^2^ = 0.25, *p* = <0.01, permutations = 999).

**Fig. 5.**
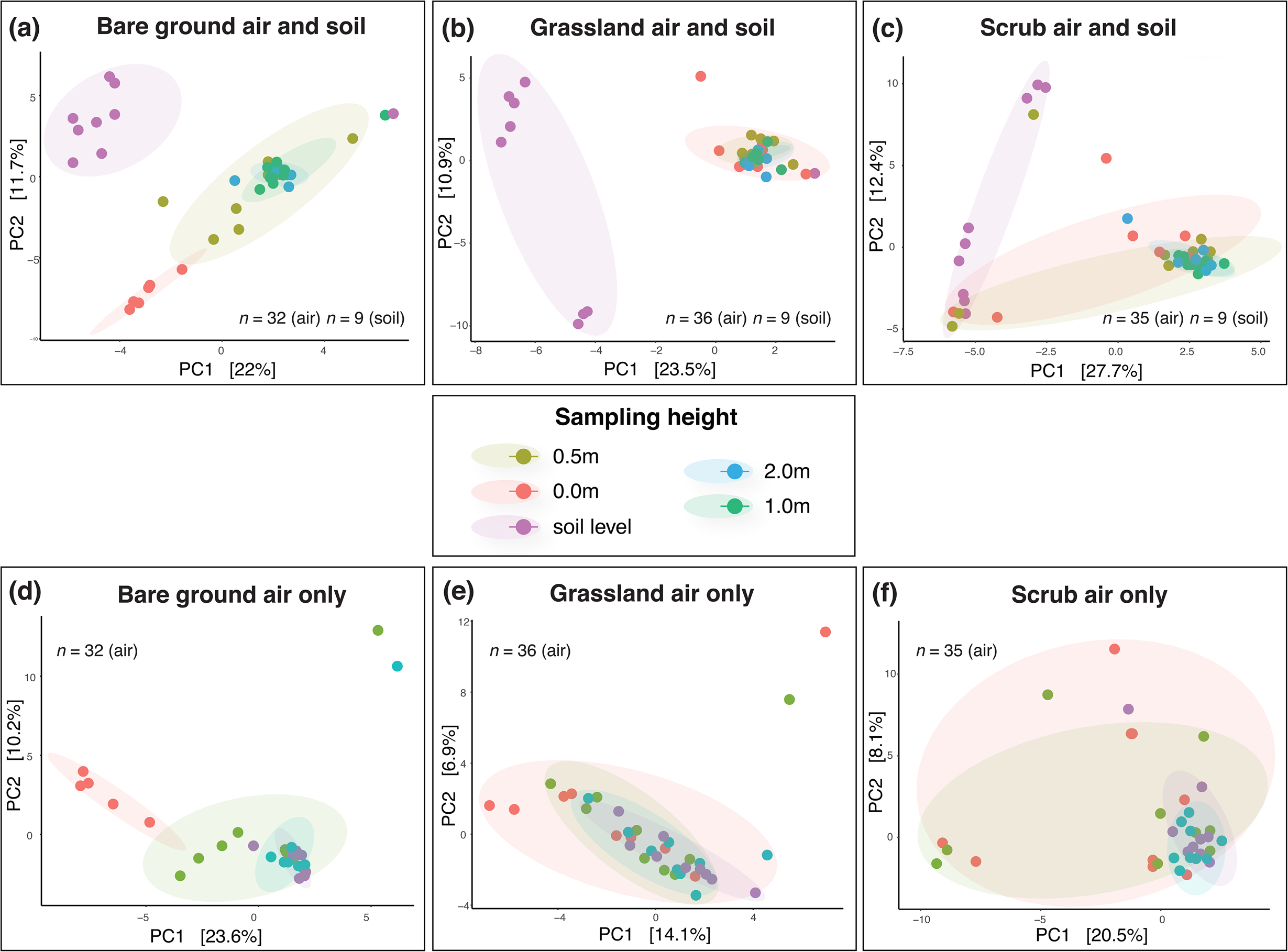
Visualising bacterial beta diversity with ordination plots of Aitchison distances based on clr-transformations of OTU abundances for each sampling height across the three habitats: (a) Bare ground air and soil, (b) Grassland air and soil, (c) Scrub air and soil, (d) Bare ground air only, (e) Grassland air only, and (f) Scrub air only. Ellipses represent Euclidian distance from the centre – with the radius equal to the confidence level (0.95). Clusters suggest differences between communities at different sampling heights (indicated by the colours).

#### *Grassland v*ertical stratification: beta diversity

Air sampling heights in the grassland habitat displayed disparate bacterial communities to the soil (**Fig. 5, b**). Sampling height explained 24% variation in bacterial community composition when all air sampling heights were included (PERMANOVA df = 4, F = 3.17, R^2^ = 0.24, *p* = <0.01, permutations = 999). However, analysis of grassland air samples in isolation showed that sampling height only explained 9% variation in bacterial community composition (**Fig. 5, e**), and was not statistically significant (df = 3, F = 1.06, R^2^ = 0.09, *p* = 0.24, permutations = 999).

#### *Scrub v*ertical stratification: beta diversity

Sampling heights in the scrub habitat displayed disparate bacterial communities (**Fig. 5, c**). Sampling height explained 22% variation in bacterial community composition when all air sampling heights and soil were included (PERMANOVA df = 4, F = 2.9, R^2^ = 0.22, *p* = <0.01, permutations = 999). Analysis of air samples in isolation showed that sampling height still explained 11% variation in bacterial community composition (**Fig. 5, f**) (df = 3, F = 1.30, R^2^ = 0.11, *p* = 0.03, permutations = 999).

#### Vertical stratification: aerobiome network analysis

In spite of differences in bacterial community composition and alpha diversity among the three study sites, network analyses showed an increase in the community complexity and interactions, defined by node degree and network size, at lower heights as compared to higher heights (**Fig. 6**). Bacterial OTUs in the scrub habitat at 0 to 0.5 m heights had the highest node degree, while the OTUs in the grassland habitat 1 to 2 m had the lowest node degree. At lower heights, the average association of any OTU in the grassland was less (node degree = 2.7) than the average association of OTUs for scrub (node degree= 4.9) and bare ground (node degree= 4.7) habitats. At upper heights, node degree for OTUs was highest for bare ground (2.7) followed by scrub (1.8) and grassland (1.7). Evaluation of link type, either positive or negative links, suggested a positive association among most OTUs, except for scrub 1 to 2 m which only had a small number of negative associations (**Fig. 6**). Comparisons of modularity between heights across the study sites suggested an increase in the network modularity at higher heights, despite the decrease in network connectance and node degree. Percentage of change in the modularity between heights was highest in the grassland (~ 50 %), although there were fewer nodes per module.

**Fig. 6.**
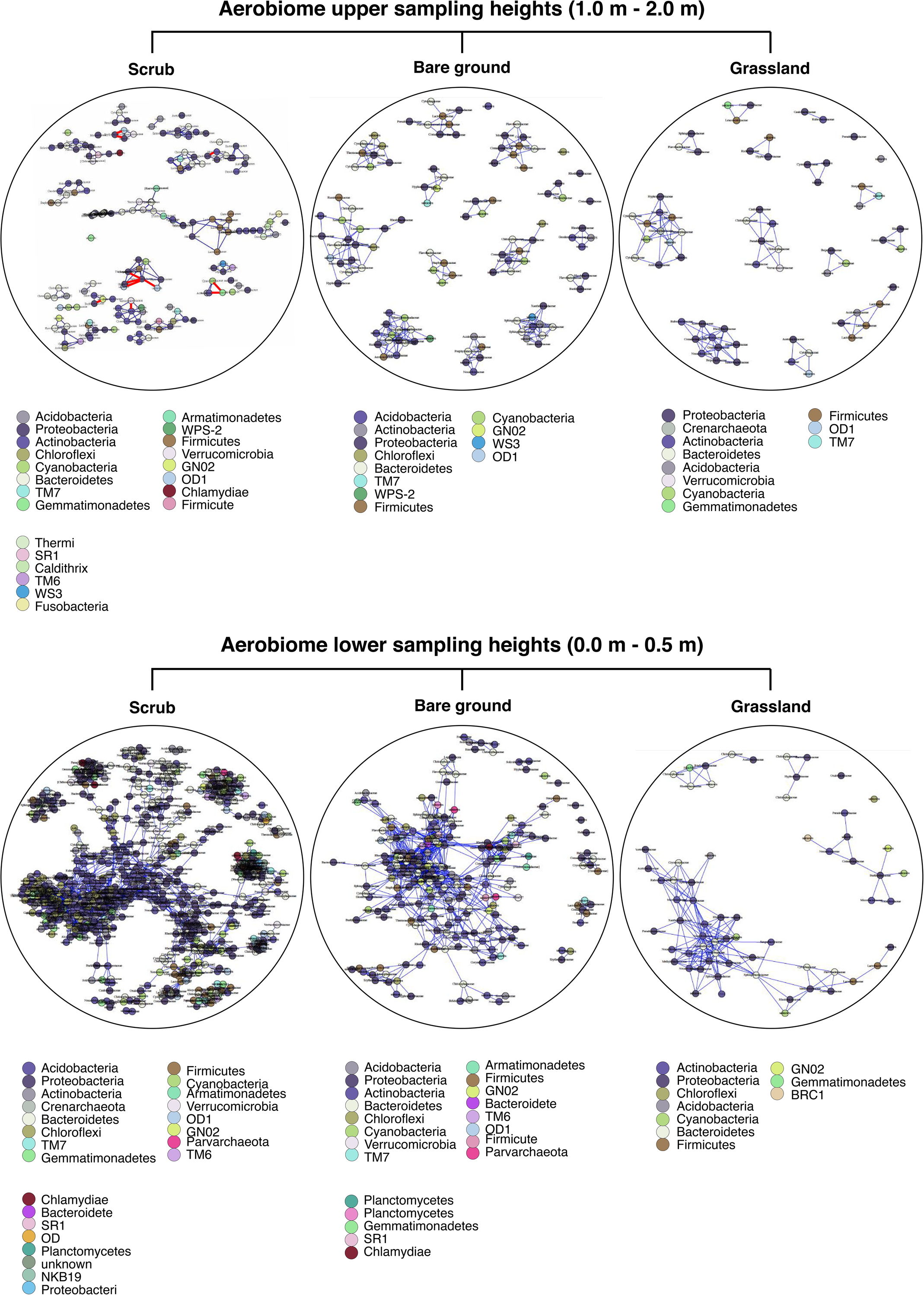
Association networks of bacterial OTUs for two vertical stratifications: 1.0-2.0 m (top panel) and 0.0-0.5 m (bottom panel). Node colour represents phylum, and node are labelled at the family level. Blue links represent positive associations, and red links represent negative associations.

### The relationship between tree metrics and bacterial alpha diversity

In the air samples, we found strong positive correlations between tree density (based on count of trees in a given radius) and bacterial alpha diversity in the 10 m radius (Spearman’s *r*_*s*_ = 0.67, *ß* = 0.67 (0.4 − 0.8), *p* = <0.01) and 25 m radius (*r*_*s*_ = 0.54, *ß* = 0.54 (0.2 − 0.7), *p* = <0.01) (**Fig. 7, a** **and** **b**). We also found significant moderate positive correlations between tree density and bacterial alpha diversity in the 50 m (Spearman’s *r*_*s*_ = 0.46, *ß* = 0.46 (0.1 − 0.7), *p* = 0.00) and 100 m radii (Spearman’s *r*_*s*_ = 0.50, *ß* = 0.50 (0.2 − 0.7), *p* = <0.01) (**Fig. 7, c** **and** **d**). Relationships between tree density and bacterial alpha diversity in soil were not statistically significant (Spearman’s *r*_*s*_ = 0.33, *p* = 0.38).

**Fig. 7.**
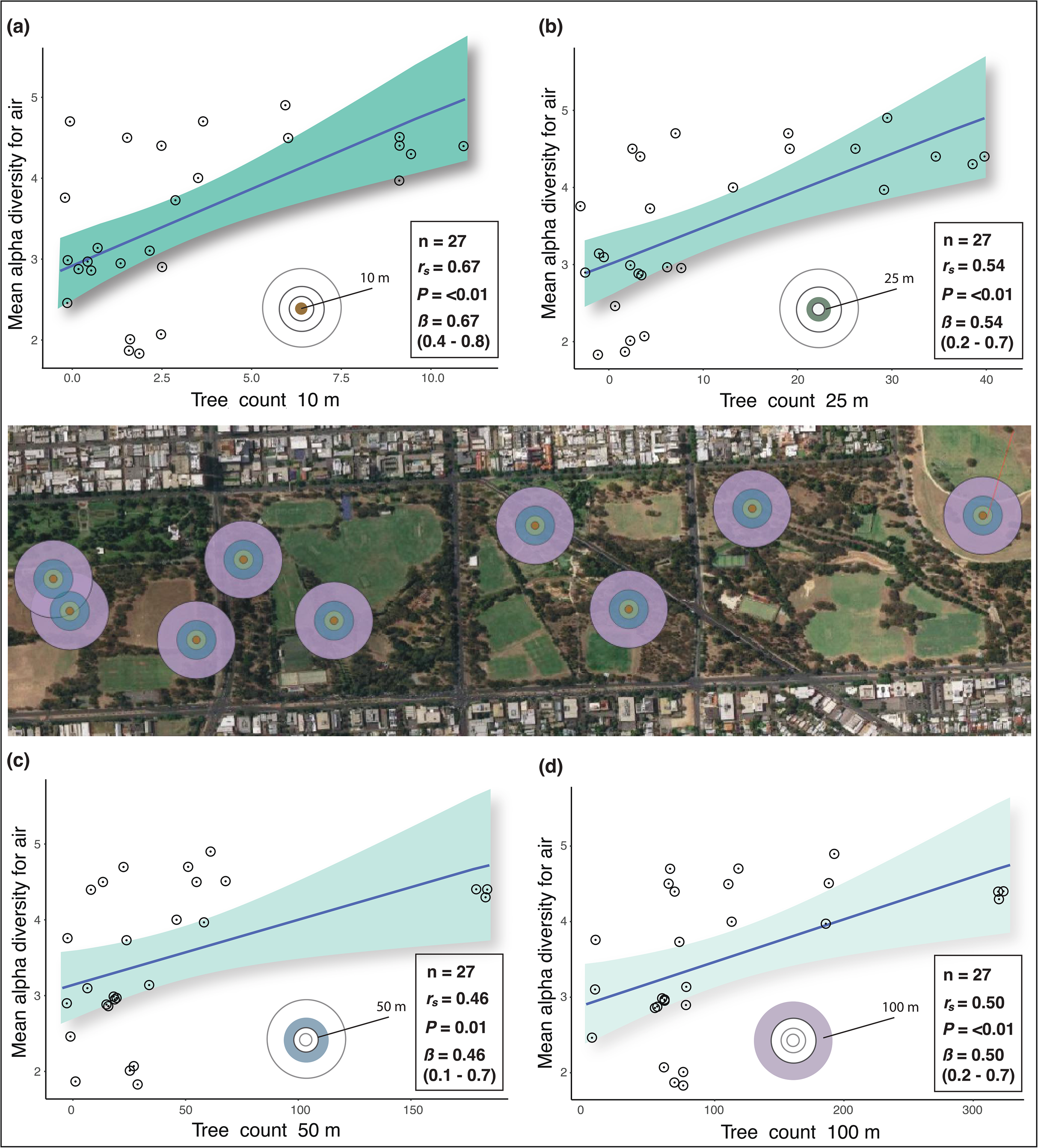
Scatterplots of Spearman’s correlations (*r*_*s*_) between bacterial alpha diversity (for all habitats and air-only samples) and tree count within each buffer radius: (a) 10 m radius from sampling points, (b) 25 m radius, (c) 50 m radius, and (d) 100 m radius. X-axis shows count of trees within buffer radii. Y-axis shows bacterial alpha diversity of air-only samples using the Shannon diversity index (H). Green shading indicates the 95% confidence intervals for each linear regression. The buffer radii are in the central aerial image and the corresponding spatial rings are in the plots. Inset also shows bootstrap results (ß) with 2.5% and 97.5% slopes.

We found significant moderate negative correlations between distance to nearest trees (from sampling stations) and aerobiome alpha diversity (Spearman’s *r*_*s*_ = −0.58, *ß* = −0.58 (−0.7 – −0.3), *p* = <0.01), and soil bacterial alpha diversity (Spearman’s *r*_*s*_ = −0.40, *ß* = −0.40 (−0.6 – −0.1), *p* = 0.03) (**Fig. 8, a** **and** **b**, respectively). Moreover, we found significant moderate positive correlations between tree canopy coverage and bacterial alpha diversity of the air in the 10 m (Spearman’s *r*_*s*_ = 0.51, ß = 0.51 (0.2 – 0.7), *p* = <0.01), 25 m (Spearman’s *r*_*s*_ = 0.66, ß = 0.66 (0.4 – 0.8), *p* = <0.01), and 100 m radii (Spearman’s *r*_*s*_ = 0.7, ß = 0.7 (0.5 – 0.8), *p* = <0.01) (**Fig. 8, c, d** **and** **f**, respectively). There was a negative correlation between canopy cover and bacterial alpha diversity in the 50 m radius that was not statistically significant (Spearman’s *r*_*s*_ = −0.27, ß = −0.27 (−0.6 – 0.2), *p* = 0.17) (**Fig. 8, e**).

**Fig. 8.**
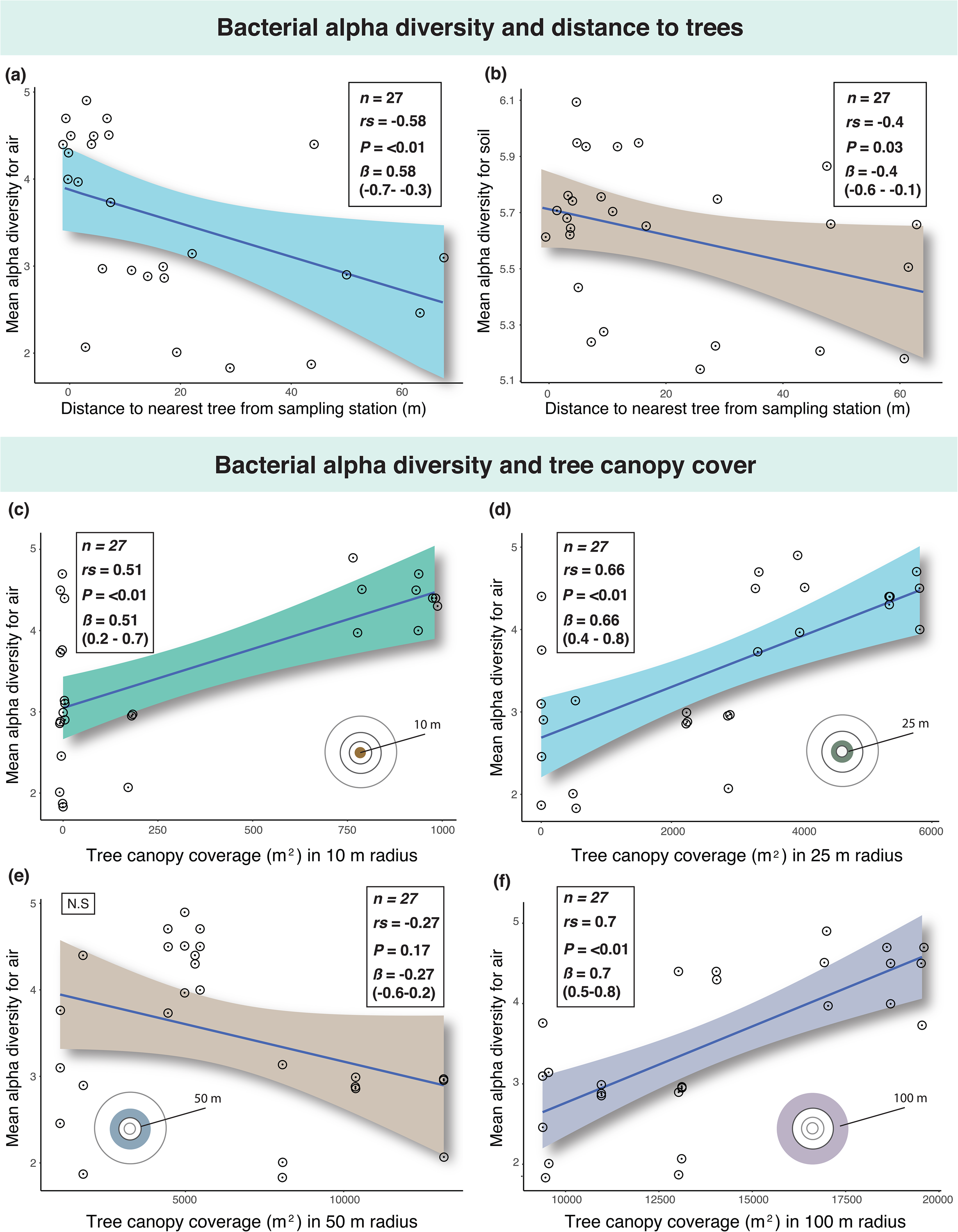
The upper panels show scatterplots of Spearman’s correlations (rs) between bacterial alpha diversity (for all habitats) and distance to nearest trees: (a) air-only samples, and (b) soil-only samples. X-axis shows distance (m) to nearest trees from sampling points. The lower panels show scatterplots of Spearman’s correlations between bacterial alpha diversity (for all habitats) and tree canopy coverage within the sampling point radii: (c) 10 m, (d) 25 m, (e) 50 m, and (f) 100 m. These relate to air-only samples. X-axis shows tree canopy coverage (m^2^). Y-axis for both upper and lower panels shows bacterial alpha diversity of samples according to the Shannon diversity index (H). Coloured shading indicates the 95% confidence intervals for each linear regression. Inset also shows bootstrap results (ß) with 2.5% and 97.5% slopes. N.S (not significant).

### Differentially abundant and notable taxa

There were 53 differentially abundant genera across habitat types (based on log‐2 fold‐change with adjusted *p* = <0.05). The top three, for example, in the scrub habitat were: *Gillisia, Sphingobium*, and *Kutzneria*; in grassland: *Parvibaculum, BSV43,* and *Pseudomonas*; and in bare ground: *Rudanella*, *Bacteroides*, and *Actinomyces*. We also observed vertical stratification of differentially abundant taxa. In the bare ground habitat, 77 genera were differentially abundant and significantly increasing in abundance with sampling height, and 97 were significantly decreasing. In the grassland habitat, 137 genera were differentially abundant and significantly increasing with sampling height, and 52 were significantly decreasing. In the scrub habitat, 41 genera were differentially abundant and significantly increasing with sampling height (**Fig. 9, a** **to** **c**), and 37 were significantly decreasing.

**Fig. 9.**
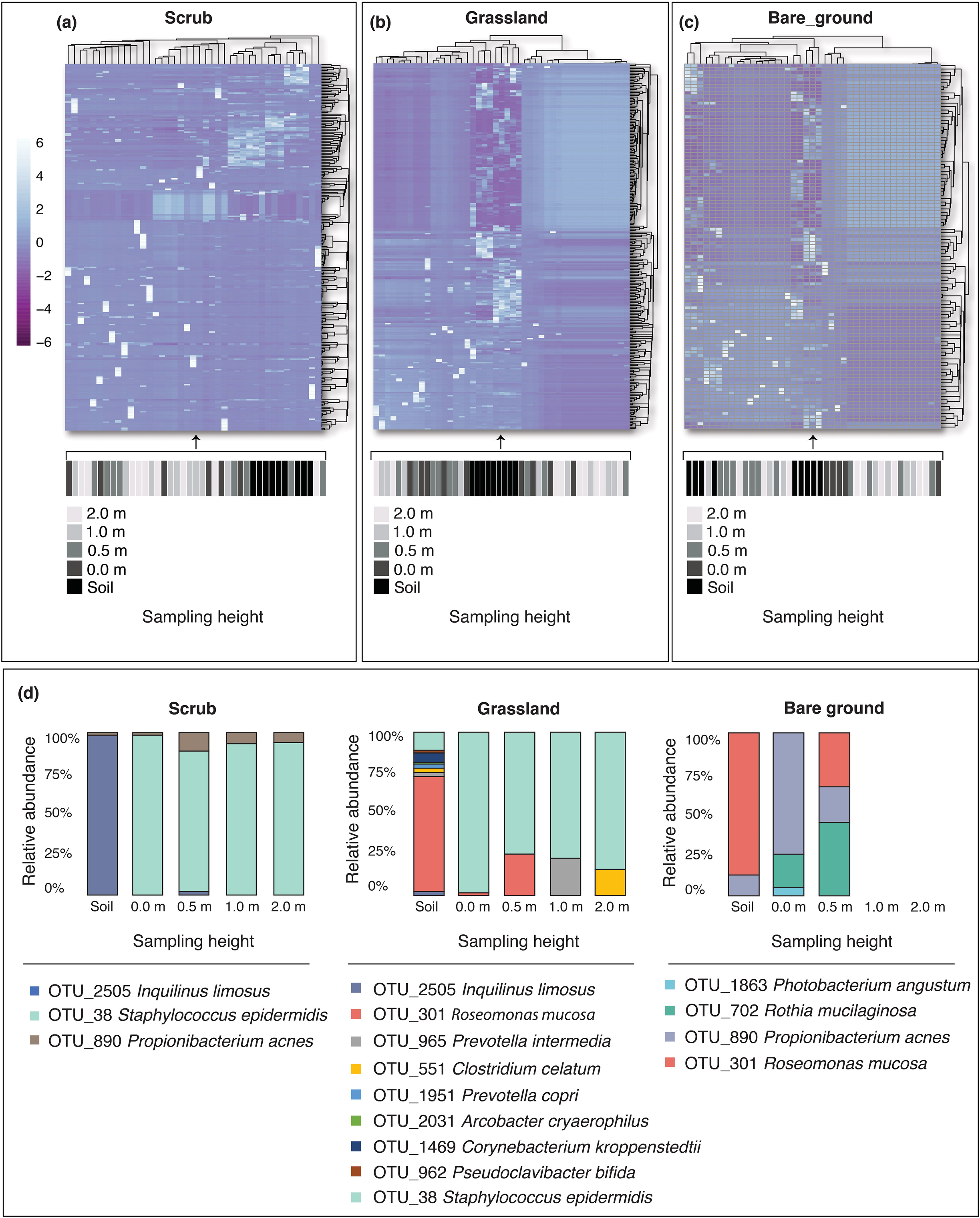
Top panels show significantly increasing (from soil level to 2 m) differentially abundant *genera* in (a) Scrub habitat, (b) Grassland habitat, and (c) Bare ground habitat measured by log‐2 fold‐change with a p value < 0.05. Extreme ends of the heat colour scale represent 6 standard deviations from the mean abundance for each genus across samples. Clustering of genera (rows) is by Manhattan distance.

We also examined differentially abundant taxa at the putative species level. After unclassified taxa were removed, we confirmed identity (100% match) via Basic Local Alignment Search Tool (BLAST) against the NCBI database (***33***). In the bare ground habitat, we found 30 differentially abundant taxa assigned at the putative species level. Sixteen of these significantly decreased in relative abundance with sampling height and 14 significantly increased (*p* = <0.01). In the grassland habitat, we found 40 differentially abundant taxa assigned at the putative species level. Thirty-two of these significantly decreased with sampling height and 8 significantly increased (*p* = <0.01). In the scrub habitat, we found 16 differentially abundant taxa assigned at the putative species level. Ten of these significantly increased with sampling height and 6 significantly decreased (*p* = <0.01). Using BLAST and a literature search, we found putative differentially abundant human pathogens in each habitat (**Fig. 9, d**). A 2-sample test for equality of proportions with continuity correction showed a significant difference in proportions of identifiable pathogenic species between grassland and scrub (Chi-squared = 5.57, df = 1, *p* = <0.02) but not between other habitats, where grassland samples exhibited significantly greater proportions of identifiable pathogenic species compared to scrub. Moreover, 87% of these significantly decreased with sampling height based on log‐2 fold‐change differential analysis (*p* = <0.01). These taxa contain bacteria that have been associated with a number of diseases, including infective endocarditis (*Rothia mucilaginosa*) and gut mucosal damage (*Prevotella copri*). More information on these diseases can be found in Supplementary Materials, Appendix C.

Shaded bars under the heatmap represent sampling heights as indicated by the corresponding colour key (where the lighter the shade, the higher the sampling height). Bottom panel (d) shows potentially pathogenic and differentially abundant *species* for each habitat and their relative abundance for each sampling height. More details on the diseases these pathogens may cause are in Supplementary Materials, Appendix C.

### Environmental metadata

In terms of the environmental metadata, there was only one significant association with bacterial alpha diversity; aerobiome alpha diversity decreased significantly in scrub habitat as windspeed increased (Spearman’s *r*_*s*_ = −0.88, *ß* = −0.88 (−0.98 – −0.5), *p* = <0.01) (full details in Supplementary Materials, Appendix D).

## DISCUSSION

Here we show that aerobiome alpha and beta diversity (community composition) differed significantly between urban green space habitat type, and that aerobiome diversity, composition and network complexity also stratified vertically. The level to which this occurred was dependent on habitat type. Therefore, potential bacterial exposure levels and transfer loads to humans will likely differ depending on habitat type as well a person’s height and behaviour. Our results confirmed that more trees, closer proximity to trees, and greater canopy coverage associate with higher aerobiome diversity, which could have important implications for landscape management and public health as growing emphasis is placed on designing and managing green spaces for wellbeing (***34***). We also found that grassland samples exhibited significantly greater proportions of identifiable pathogenic bacteria compared to scrub, and their abundance decreased significantly with sampling height. Our study was conducted only in the Adelaide Parklands, South Australia and therefore may not be representative of urban green spaces in other areas. Future work should explore these trends in additional geographical, socioeconomic, cultural areas to understand both generalisability and opportunities to optimise green space exposure for health benefits.

### Aerobiome compositional differences between habitats

The scrub habitat exhibited the most biodiverse aerobiome in our study. This corroborates other studies that suggest that environmental microbiomes are more biodiverse in urban habitats with more complex vegetation communities (***35, 36***). Growing evidence suggests that exposure to biodiverse environmental microbiomes could have important implications for human health (***5, 14, 37***). For example, environmental microbiomes are essential in the development and regulation of immunity (***1, 6**),* and soil-derived butyrate-producing bacteria may supplement gut bacteria and have anxiety-reducing effects (***5***). Importantly, urban green space exposure can result in transmission of environmentally-derived bacteria to the skin and airways (***21***). Furthermore, a recent study showed that transfer of bacteria from biodiverse environments enhanced immunoregulatory pathways in children (***38***). Consequently, environments with different levels of bacterial diversity may affect the potentiality of bacterial exposure levels and transfer loads, warranting further research. We found differentially abundant putative pathogenic taxa and showed significant differences in proportions between grassland and scrub habitat samples. In other words, amenity grassland seemed to exhibit a significantly greater proportion of (identifiable) pathogenic species compared to scrub samples. However, considerably more research is needed to fully explore the validity and generalisability of these results. As with many microbial ecology studies, only identifiable bacterial taxa were used in the differential abundance and analyses that identified the pathogenic taxa (i.e., unclassified taxa were removed). This could result in recording bias with implications for validity.

Our results suggest that tree density, distance to nearest trees, and tree canopy cover could have a considerable influence on aerobiome alpha diversity. This corroborates reports of trees acting as stationary vectors, spreading bacterial cells in the air (***39***). Complex plant detritus (leaf litter) and organic matter at the base of trees, and corresponding soil-microbe systems, may also contribute to tree-associated aerobiomes. The number of trees and amount of canopy coverage within a given radius correlated strongly with alpha diversity. Furthermore, negative correlations were shown between distance to nearest trees and bacterial alpha diversity for air and soil. This supports the results of the tree density associations and suggests that closeness to trees could be important. These results could have important implications for landscape management and public health. Indeed, there have been widespread calls to improve urban ecosystem services by augmenting tree coverage (e.g., to help reduce urban heat island effects (***40***), support wildlife (***41, 42***), improve sleep (***43, 44***), and capture precipitation to reduce flood risk (***45***)). There is also a need to restore complex vegetation communities and host-microbiota interactions that provide multifunctional roles in urban ecosystems (***37, 46, 47***). An important limitation in our study was that tree species and structural diversity metrics were not used. These additional measures could have enriched the quality of analysis and implications of our results and further research that takes these factors into account is needed. However, our findings suggest additional co-benefits from increasing urban tree coverage due to its potential to mediate aerobiome alpha diversity. Our results also corroborate other studies showing microbial alpha diversity increasing along densely-urban to semi-natural environmental gradients (***48, 49***).

Our results suggest that aerobiome beta diversity (compositional differences) differs between habitats. The results imply that microbial communities in the soil of the scrub habitat are significantly different to bare ground and grassland, which are more compositionally aligned. It is possible that bacterial homogeneity between grassland and bare ground is attributed to homogeneity of vegetation complexity (***50***). In other words, phyllosphere (total above-ground portion of plants) and rhizosphere (soil root zone) presence and complexity create conditions for different microbial relationships and thus compositional disparity with less botanically-complex or depauperate habitats (***36, 37***). Taken together with the alpha diversity results, significantly more bacterial species and unique communities exists in scrub habitat samples compared to grassland and bare ground samples. This could mean that humans are exposed to a greater diversity of bacteria in the scrub habitat. Future studies should focus on the functional relevance of these findings.

### Aerobiome vertical stratification

In our study, vertical stratification in bacterial alpha and beta diversity occurred in the bare ground and scrub habitat. However, for the grassland aerobiome, both alpha and beta diversity were relatively stable as height increased. This is the first study to demonstrate that aerobiome vertical stratification is contingent on habitat type, which is important for potential human exposure. As mentioned, urban green space exposure can result in transfer of environmental bacteria to the skin and respiratory tract (***21***), and our study shows that the composition and diversity of aerobiome bacteria may differ between heights (from ground level to 2 m). Consequently, there could be different bacterial exposure levels and transfer loads depending on a person’s height and activity (***27***), however, further confirmatory research is needed. Our results suggest that this may not be the case in amenity grassland where bacterial alpha and beta diversity exhibited high levels of homogeneity among heights. Further research is required to determine the reasons for the lack of vertical stratification in grassland. However, we hypothesise that lower baseline diversity, bacterial resources, openness and airflow in this habitat may be contributing factors. Our study also provides some evidence that different urban green space habitats and heights may not only affect exposure levels and transfer loads of bacterial diversity, but also the presence of notable and potentially pathogenic species for humans. The relative abundance of pathogens identified in the grassland habitat decreased significantly with sampling height. It is possible that a number of these potential pathogens may originate from larger air-sheds (consistent with increasing relative abundance with height), however grasslands may have lesser capacity, compared to scrub or bare ground, to present barriers to this broader airflow or contribute to a more locally distinctive aerobiome. These findings highlight the need for further empirical studies focusing on functional interactions in the environment-aerobiome-health axis.

Our network analyses also provided evidence to support aerobiome vertical stratification. We saw a decrease in bacterial interactions and network complexities with increased network modularity at higher heights compared to lower heights across habitats, which might be attributed to reduced bacterial diversity with sampling height. This pattern might be due to increasing influence, with increasing height, of diluted and somewhat homogenised aerobiomes from larger airsheds, representing the physical mixing of air (and therefore aerobiomes) from multiple different and distant ecological sources. Increased modularity with reduced network size and interactions may also indicate the existence of relatively simplified, yet modular bacterial communities at higher heights. This could be the function of sparse food resources, especially if associations in the networks reflect niche-based interactions. Increased modularity indicates the presence of dense connections between bacteria within modules but sparse connections between bacteria in different modules, whereas reduced connectance means reduced probability of interactions between any pair of bacteria. Increased modularity with reduced connectance often indicates ecological stability (***51***). Moreover, presence of mostly positive associations might also suggest cooperation for resources or lack of competition among the interacting OTUs in the community. While association-based networks allow a depiction of potential interactions among OTUs and portray community structure, they do not separate niche-based and biological interactions. Experiments with cultures are recommended to dissociate interaction types and understand the biological and ecological mechanisms behind the observed interactions and network complexity. This action could be important to gain a greater ecological understanding of aerobiome assembly (including vertical stratification), dynamics, and the potentiality of bacterial exposure. Our results provide strong evidence that vertical stratification is a key factor not only in aerobiome diversity and composition, but also in aerobiome interactions, community structure and complexity.

In conclusion, our study provides evidence that bacterial alpha and beta diversity differed significantly between habitats, with scrub habitat providing the most biodiverse aerobiomes. We provide evidence supporting the presence of aerobiome vertical stratification in bacterial community diversity, composition and complexity, which also differed in a habitat-dependent manner. Our results confirmed that more trees, closer proximity to trees, and greater canopy coverage associated with higher alpha diversity of the aerobiome. Finally, we found that grassland samples exhibited significantly greater proportions of identifiable putative pathogenic bacteria compared to scrub, and their richness decreased significantly with sampling height. As discussed, there is growing evidence to suggest that exposure to biodiverse aerobiomes may contribute towards the development and regulation of immunity and support mental health (***1, 4–6***). Gaining a greater understanding of bacterial transmission routes, exposure levels, transfer loads, and downstream health implications is required. This aerobiome characterisation study provides novel insights into the urban ecosystem to help encourage further empirical investigations. Future research should focus on the functional interactions between humans and the aerobiome. Although additional research is required, our findings also support calls to increase urban tree cover. Exploring the mediatory roles of trees in aerobiome compositional and functional diversity could have important implications for landscape management and public health.

## MATERIALS AND METHODS

### Site selection

Our study was undertaken in the southern Adelaide Parklands (Kaurna Warra Pintyanthi), South Australia, which comprised nine vegetated plots that spanned approx. 18 ha. The nine plots included three amenity grasslands, three scrub, and three bare ground (exposed soil) habitats.

There were several justifications for selecting this site: (1) the southern Parklands occur within the Upper Outwash Plain soil boundary (coalescing alluvial soil, draining the Eden Fault Block), which provided broad consistency in soil geochemistry; (2) a single section of the Parklands provided control over potential micro-geographic variation effects on the aerobiome (e.g., distance to coast, elevation, orientation, aspect, and dominant vegetation communities); (3) the Parkland habitats are representative of the types of green spaces that urban residents are regularly exposed to when commuting or recreating; and, (4) the City of Adelaide provided guidance in the selection process, identifying accessible (and inaccessible) plots.

We defined the boundaries of the nine plots (as polygons) in QGIS 3 (v3.0.2) in conjunction with the City of Adelaide. Using spatial shapefiles for the plot boundaries, we generated random point algorithms to provide random sampling points within each of the nine study plots (**Fig. 10, a**). We recorded geographic coordinates for each sampling point and programmed them into a handheld global positioning system (GPS) receiver. We operated the GPS receiver in the field, allowing us to pinpoint the locations for the sampling stations.

**Fig. 10.**
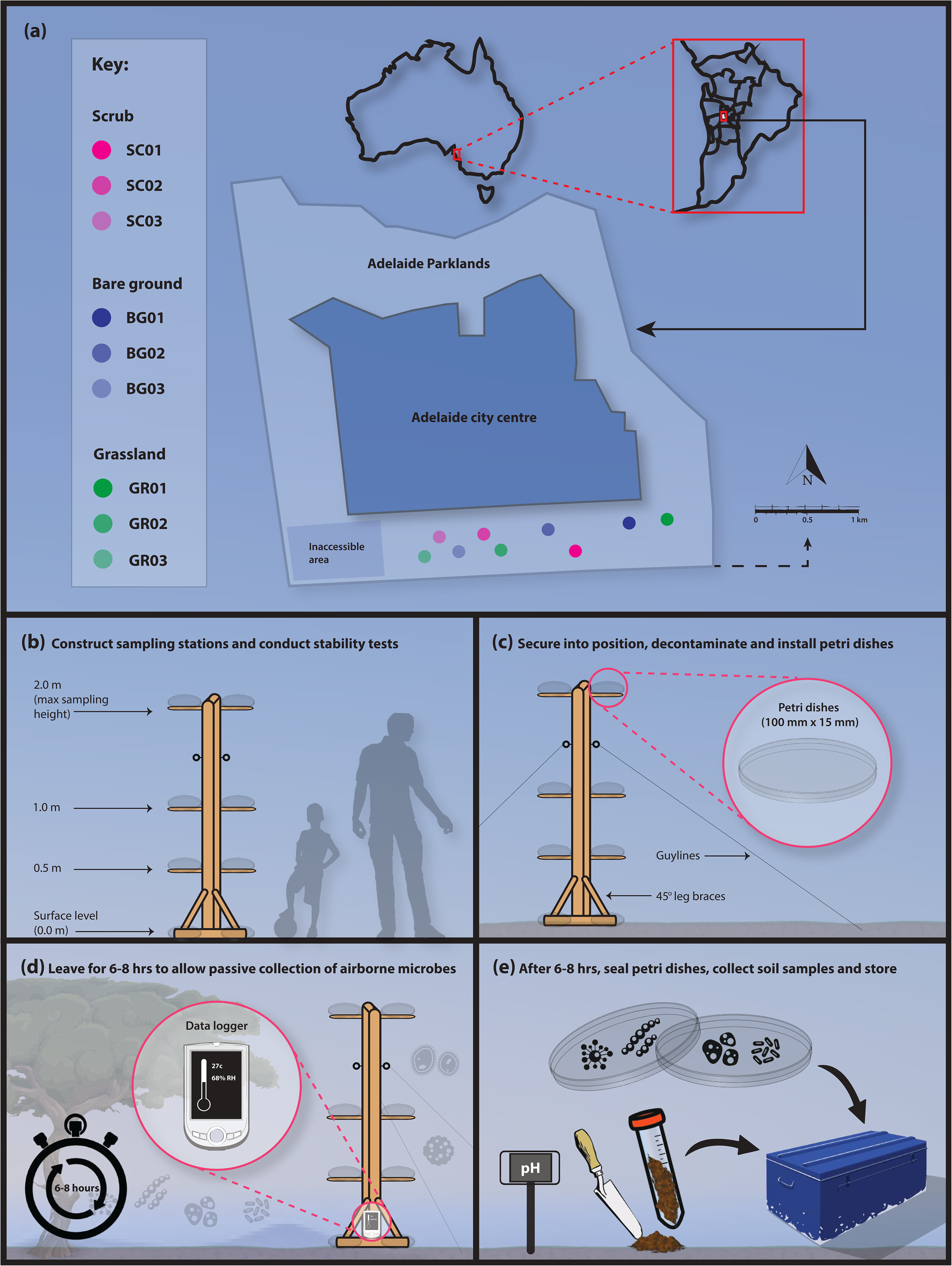
(a) Location of study sites, showing the different habitat types and randomly selected sampling locations. (b-e) Vertical stratification sampling station and methods used to collect (passively) air and (actively) soilborne bacteria. We installed the stations in three different habitat types in the Adelaide Parklands.

### Sampling equipment

Sampling stations (**Fig. 10, b** **to** **e**) were constructed using timber (42 mm × 28 mm × m), steel brackets, hooks and guy-lines (***27***). We secured lab-grade clear plastic petri dishes (bases and lids) to the sampling stations, which were used to sample the aerobiome via passive sampling (***22***).

### On-site setup

We installed the sampling stations on site between 0600-0800hrs on the 4th, 5th and 6th November 2019. At 0800hrs, sampling stations were decontaminated using a 5% Decon 90 solution. We then installed the petri dishes for passive sampling. The data loggers were also decontaminated. In the scrub habitat (defined as vegetation dominated by locally native shrubs, usually ≤5 m tall, with scattered trees) (***52***), the nearest trees and shrubs were between 2-5 m from the sampling stations, and all trees were <10 m height and 20-50 cm in diameter at breast height (***25***).

### Sampling protocol

We installed temperature and relative humidity data loggers at each sampling station (***27***). We programmed each logger to record data at 8-second intervals for the entire sampling period. At the start of each sampling day, we calibrated the dataloggers using a mercury thermometer (Gerotherm) and a sling psychrometer (Sper Scientific 736700). We collected other metadata including windspeed and soil pH (Alotpower digital meter). We inserted the pH meter into the soil for a period of 1-minute before taking a reading (manufacturer’s instructions). We obtained data for windspeed and direction from Adelaide’s meteorological weather station at Ngayirdapira (West Terrace): Lat: −34.93, Lon: 138.58, Height: 29.32 m. We also used a handheld anemometer (Digitech *QM-1644)* to record these parameters hourly at each sampling site (***22***).

### Soil samples

We used a small, decontaminated shovel to collect topsoil samples and stored these in sterile 50 mL falcon tubes. We collected five topsoil samples (approx. 0-5 cm depth) at equidistant sampling points, 20-30 cm from the stem of each sampling station (***53***). We pooled and homogenised the soil samples, passed them through a decontaminated 1 mm pore sieve, and placed them in new sterile and labelled 50 mL Falcon tubes. We included field controls for the soil by opening sterile falcon tubes for 60 s in the equipment box at each site (***54***). We placed all soil and field control samples immediately into an ice box, and stored the samples in the lab at −80°C prior to further processing (***55***). In total, we collected 45 soil subsamples per sampling day across nine sampling stations with three temporal replicates (over three days). We pooled and homogenised subsamples by sampling station and day, which gave a total of 27 homogenised samples (nine per sampling station) plus 9 field controls.

### Aerobiome samples

To collect aerobiome samples, we used a passive sampling technique, following established protocols (***22, 26***). We installed petri dishes (100 mm × 15 mm) with Velcro tabs on the sampling stations at four sampling heights: ground level (i.e., 0.0 m), 0.5 m, 1 m, and 2 m. The total height of the sampling station was 2 m from ground level (95% of typical adult male heights lie within 2 SD at 1.93 m, and 1.78 m for females based on a study across Europe, North America, Australia and East-Asia) (***56***). Various human characteristics informed the height selection (e.g., representation of adults vs. children, and different activities such as sitting, crawling, walking) (***57, 58, 27***). We decontaminated the steel plates supporting the petri dishes with 5% Decon 90 solution.

We secured the petri dishes to the sampling stations (**Fig. 10**), leaving them open for 6-8 hours (***22***), and closing them at the end of the sampling period. To reduce contamination, new disposable laboratory gloves were worn for each vertical sampling point. Once sampling was complete, we sealed the petri dishes using Parafilm, labelled and transported them to the laboratory (on ice) for storage at −80°C (***26***). We collected field control samples by leaving unused petri dishes for 60 s in the equipment box and sealing them at each site.

### DNA extraction, amplification and sequencing

We extracted DNA from soil and air samples at the facilities of the Evolutionary Biology Unit, South Australian Museum. Using a digital number randomiser, we processed samples on a randomised basis. We processed the low biomass air samples prior to the higher biomass soil samples to minimise cross-contamination.

To extract DNA, we swabbed the petri dishes in the lab using nylon-flocked swabs (FLOQSwabs Cat. No. 501CS01, Copan Diagnostics Inc.) (***5, 26, 27, 59***). All swabbing was carried out in a laminar flow cabinet type 1 (License No. 926207) and each sample was swabbed for 30 s. Samples from the base and lids of each petri dish for each height, station and date were pooled. We cut the swabs directly into Eppendorf tubes (2 mL). We used Qiagen QIAamp DNA Blood Mini Kits to extract DNA from the swabs and extraction blank controls. For extraction blank controls, we used sterile water and reagents instead of a sample and all DNA extraction steps were performed as if they were normal samples. To extract DNA from the soil samples (and extraction blank controls), we used Qiagen DNAeasy PowerLyzer Soil Kits and followed the manufacturer’s instructions. PCR amplification was conducted in triplicate using the 341F/806R primer targeting the V3-V4 region of the 16S rRNA gene (5’ -CCTAYGGGRBGCASCAG-3’/5’ -GGACTACNNGGGTATCTAAT-3’). The 300 bp paired end run was sequenced on an Illumina MiSeq platform at the Australian Genome Research Facility using two flowcells (ID 000000000-CW9V6 and 000000000-CVPGT). We conducted image analysis in real-time by the MiSeq Control Software (v2.6.2.1) and Real Time Analysis (v1.18.54). We used the Illumina bcl2fastq (2.20.0.422) pipeline to generate sequence data.

### Bioinformatics and statistical analysis

Raw 16S rRNA gene sequences processing, OTU picking, taxonomic assignments, and decontamination were as per Robinson et al. (2020) (described in detail in Supplementary Materials, Appendix A). To estimate OTU alpha diversity we derived Shannon Index values (***5***) in phyloseq (***60***) in R. Prior to analysis of compositional data, we used centre log-ratio (clr) transformations (***61***). Information acquired from this approach is directly relatable to the environment (***62***). We generated violin plots with ggplot2 (***63***) to visualise the distribution of the alpha diversity scores for each habitat and height. Bacterial beta diversity was visualised using ordination plots of Aitchison distances based on clr-transformations of OTU abundances. Ordination plots show low-dimensional ordination space in which similar samples are plotted close together, and dissimilar samples are plotted far apart.

We used permutational multivariate analysis of variance (PERMANOVA) to test for compositional differences between different sites, habitats and sampling heights, and permutation tests for homogeneity of multivariate dispersions using vegan (***64***) in R. Pearson's product-moment and Spearman’s rank correlation tests were used to examine correlations between habitat, sampling height and alpha diversity scores. Using phyloseq, we calculated OTU relative abundances to examine the distribution of taxa that may have potential implications for public health. We used DESeq2 (***65***) in R to conduct differential abundance analysis based on log‐2 fold‐change. To compare presence and proportions of taxa we used 2-sample tests for equality of proportions with continuity corrections. We also applied bootstrap resampling to assign a measure of accuracy to sample estimates for the Spearman’s correlations, using a minimum of 1,000 iterations. This was carried out with the psych (***66***) and boot (***67***) packages in R.

In order to understand the effect of vertical stratification on bacterial interactions and community structures, we evaluated association-networks of bacterial OTUs. We combined the OTU database from 0-0.5 m and 1-2 m for each site, and constructed two networks per site (i.e., lower and upper height), such that in total six networks were evaluated across the three habitats. In the evaluated network, nodes represent OTUs and links exists between a pair of OTUs if their frequencies are significantly associated (absolute abundance > 0.7, *p* = < 0.01). The type of association, whether positive or negative, was represented with blue and red links, respectively. To account for compositional bias associated with OTU data, we used SparCC (***68***) to define associations, and only OTUs with sequence counts >10 were included. Randomly permuted (*n =* 100) data were used to estimate significance of associations, and igraph (***69***) was used to visualize and evaluate the plots. We also ran Spearman’s correlation tests with bootstrap resampling to determine whether environmental metadata (pH, temperature, windspeed) associated with bacterial alpha diversity. Outliers were considered as data points more than 1.5 × above the third quartile or below the first quartile.

### Geospatial analyses

We investigated possible relationships between aerobiome samples and surrounding vegetation properties using spatial buffer zones. For the buffer analysis, we used vector geoprocessing tools in QGIS 3. Buffer sizes of 10 m, 25 m, 50 m, and 100 m were considered appropriate for the study scale. Similar distances have been used in previous green space and epidemiology studies (***31, 70–72***). A 100 m maximum buffer radius was chosen; at greater distances, effects would no longer be local to the sampling points (i.e., they would overlap with other sampling points). To determine tree canopy cover within each buffer radii, ESRI shapefiles were imported into i-Tree Canopy (***73***). This enabled random sampling points (between 50-250 points per buffer) and selection of land cover classification and associated metrics overlaid with Landsat 8 satellite imagery (***73–75***). Tree count and distance measures were acquired using geometry tools in QGIS 3.

## Acknowledgements

We acknowledge and pay our respects to the Kaurna people, the traditional custodians whose ancestral lands we conducted the research on for this study. We would also like to acknowledge the City of Adelaide who helped to facilitate this study.

## Author contributions

J.M.R and M.F.B contributed to the conception and design of the study; J.M.R and C.C.D conducted the field and lab work; J.M.R, R.A, R.P, and C.L conducted the bioinformatics and data analysis; J.M.R wrote the manuscript; J.M.R produced the figures and data visualisations; J.M.R, C.C.D, R.A, R.C, R.P, C.L, P.W, and M.F.B contributed to manuscript internal critical review process and revisions. All authors read and approved the submitted version.

## Data and materials availability

All data and code used in this study are available on the *UK Data Service ReShare*.

